# Centrifuge+: improving metagenomic analysis upon Centrifuge

**DOI:** 10.1101/2023.02.27.530134

**Authors:** Junfeng Liu, Yunran Ma, Yong Ren, Hao Guo

**Author notes:** The authors wish it to be known that, in their opinion, the first two authors should be regarded as Joint First Authors.

## Abstract

**Summary:** Accurate abundance estimation of species is essential for metagenomic analysis. Although many methods have been developed for classification of metagenomic data and abundance estimation of species, the abundance estimation of species remains challenging due to the ambiguous reads that align equally well to more than one genome. Here, we present Centrifuge+, which introduces unique mapping rate to describe the influence of similarities among species in the reference database when analyzing ambiguous reads. In contrast to the popular Centrifuge, Centrifuge+ improved the accuracy of abundance estimation on simulated reads from 4278 complete prokaryotic genomes.

**Availability and implementation:** The source code is available at https://github.com/mNGSmethods/Centrifugep.

**Supplementary information:** Supplementary data are available at *Bioinformatics* online.

## 1 Introduction

Metagenomic sequencing has provided great improvements in microbiome analysis by metagenomic experiments that can be broadly categorized as either microbiome experiments or pathogen identification experiments (Knight *et al*., 2018; Lu *et al*., 2022). In microbiome experiments, researchers focus on describing what is present in a given sample. For pathogen identification experiments, the focus of researcher is identifying one or few pathogenic microbes. In order to achieve the goal of metagenomic experiments, estimating the abundance of the species in a given sample becomes very important in metagenomic analysis. However, the ambiguous reads that align equally well to more than one genome make challenge for the abundance estimation of the species because it is very difficult to identify the taxon of ambiguous reads. There are two reasons for causing the ambiguous reads. The first reason is that closely related species are present in a given sample. The second reason is because of the nearly identical genomes in a reference database that is used for identifying the taxon of each read. In order to overcome the challenge of ambiguous reads, a separate abundance estimation algorithm is necessary for most metagenomic classification tools. To counter the ambiguous reads caused by closely related species in the same sample, Kim et al. (2016) defined a statistical model in the popular metagenomic classification tool Centrifuge (Kim *et al*., 2016) and used it to estiamte the abundance of species through an Expectation-Maximization (EM) algorithm. In the statistical model of Centrifuge, the probability is only depended on the abundance of species *j* and the length of the genomes of species *j* when the ambiguous read *i* is classified to species *j*. However, for the ambiguous reads caused by the nearly identical genomes in a reference database, the probability is also decided by the similarities between species *j* and the other species in the reference database if the ambiguous read *i* is classified to species *j*. Although the similarities among species in the reference database have been considered in the statistical model of Bracken (Lu *et al*., 2017), which was developed to estimate species abundance in conjunction with Kraken (Wood and Salzberg, 2014), the statistical model of Bracken can be only used to analyze Kraken classification results and requires generating simulation data to estimate species abundance.

To address the above limitation, we introduce Centrifuge+, which modified the statistical model of Centrifuge and improved metagenomic analysis. In the modified statistical model, the influence of similarities among species in the reference database is described by unique mapping rate when analyzing the ambiguous reads. In addition, we use the modified statistical model to estimate species abundance through an Expectation-Maximization (EM) algorithm.

## 2 Implementation

Centrifuge+ is based on Centrifuge with the same methods of reference database sequence compression and classification of microbial sequences, but is different from Centrifuge on the statistical model, which considers the influence of similarities among species in the reference database on estimating species abundance. In order to implement the modified statistical model, we modified Centrifuge developed by Kim et al. (2016) under the terms of the GNU General Public License and named it as Centrifuge+.

### 2.1 The modified statistical model

Similar to Centrifuge, the likelihood function is defined as follows:

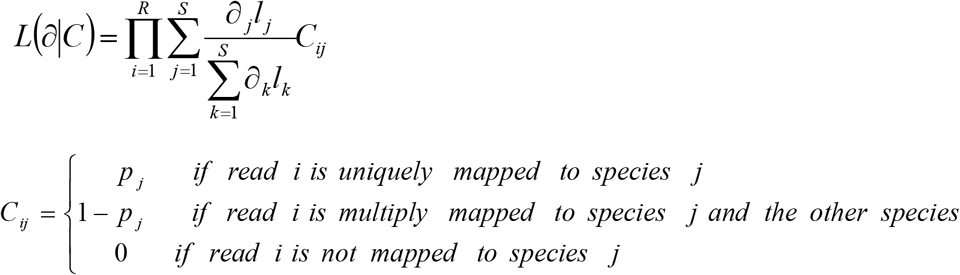

where *R* is the number of the reads, *S* is the number of species, ∂_*j*_ is the abundance of species *j* and 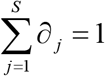, *l*_*j*_ is the average length of genomes of species *j*, and *pj* is the unique mapping rate of species *j*. For species *j*, we count the number of reads that are uniquely mapped to it, *m*. If the number of reads that can be classified to species *j* is *n*, the unique mapping rate of species *j* is *m/n*.

In the modified statistical model, we introduced the unique mapping rate (*pj*) to describe the influence of similarities between species *j* and the other species in the reference database when assigning a value to *C*_*ij*_. However, in the statistical model of Centrifuge, the value of *C*_*ij*_ is only 1 or 0 according to whether read *i* is mapped to species *j*.

### 2.2 Abundance analysis

To estimate species abundance, the following EM procedure is implemented in Centrifuge+.

Initialization step (I-step): the initial value of ∂_*j*_ is *1/S*.

Expectation step (E-step):

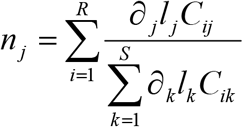

where *n*_*j*_ is the estimated number of reads mapped to species *j*.

Maximization step (M-step):

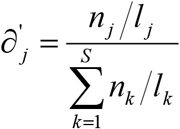

where ∂^’^_*j*_ is the updated estiamtion of species *j’s* abundance and used in the next iteration as ∂_*j*_.

Centrifuge+ repeats E-step and M-step until. 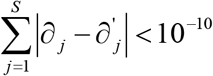. The above EM procedure is also implemented in Centrifuge except for E-step.

## 3 Results and discussion

We compared Centrifuge+ and Centrifuge by assessing the match between the estimated abundance and the true abundance distribution of genomes in the simulated reads at the species level. The simulated read data set was created from the 4278 complete prokaryotic genomes in RefSeq (Pruitt *et al*., 2014) by Kim et al. (2016). They used the Mason simulator (Luke *et al*., 2005) to generate 10 million 100-bp reads and the resulting file was named bacteria_sim10M.fa (Kim *et al*., 2016). Then, they randomly down-sampled the datasets to 10 thousand reads (bacteria_sim10K.fa) without replacement. We used this dataset for the performance comparison of Centrifuge+ and Centrifuge (Supplementary Materials). Pearson’s correlation coefficient between the true abundance and the estimated abundance of Centrifuge+ was 0.6 at the species level based on 10 thousand simulated reads (Fig. 1A). However, Pearson’s correlation coefficient was only 0.26 at the species level when comparing the true abundance and the estimated abundance of Centrifuge (Fig. 1B). If the top five percent worst abundance estimates were omitted, the correlation coefficient of Centrifuge+ can improve to 0.95 (Fig. 1C). But, the correlation coefficient of Centrifuge only can improve to 0.46 when omitting the top five percent worst abundance estimates (Fig. 1D). The above results show that the abundance estimates of Centrifuge+ are more closely matched to the true abundance than Centrifuge. Even for the top five percent worst abundance estimates, Centrifuge+ is still significantly better than Centrifuge (Supplementary Fig. S1). Moreover, the more accurate abundance estimates make Centrifuge+ to have a higher recall (75.86% VS 65.51%), which is the proportion of true positive species divided by the number of distinct species actually in the sample (Supplementary Table S1). Though Centrifuge+ was a little lower than Centrifuge on the the precision (98.33% VS 100%), which is the proportion of true positive species identified in the sample divided by the number of total identified species, Centrifuge+ has a higer F1 score (0.86 VS 0.79) that is the harmonic mean of recall and precision (Supplementary Table S1).

**Fig. 1.**
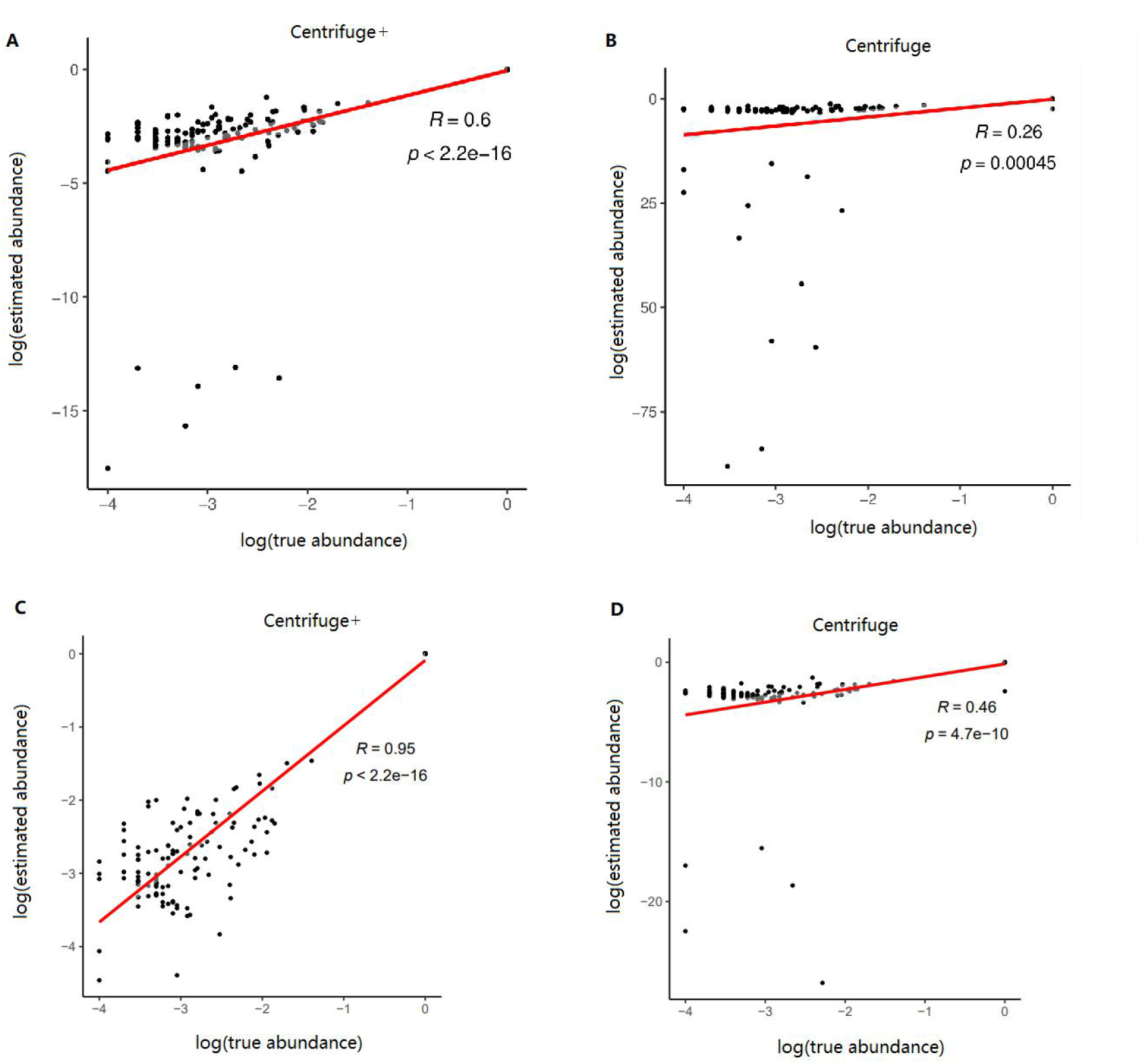
Comparison of log-scaled true abundance and estimated abundance at species level based on 10 thousand simulated reads. *R* and *p* are Pearson’ s correlation coefficient and *p*-value, respectively. (A) Comparison of log-scaled true abundance and Centrifuge+ abundance estimates; (B) Comparison of log-scaled true abundance and Centrifuge abundance estimates; (C) Comparison of log-scaled true abundance and Centrifuge+ abundance estimates when the five percent worst abundance estimates of Centrifuge+ were omitted; (D) Comparison of log-scaled true abundance and Centrifuge abundance estimates when the five percent worst abundance estimates of Centrifuge were omitted.

When describing the probability of observed read, the statistical model in Centrifuge does not distinguish between species in the processing of unique and multiple mapping reads, that is, no matter to which species the read is classified to, the unique mapping rates of different species are the same. Centrifuge’s above processing method implies the following assumption: the probability of the occurrence of unique and multiple mapping reads of different species is only determined by species abundance. However, due to the influence of reference genome similarity, the probability of unique mapping reads and multiple mapping reads of different species will also be different. For example, an observation sample contains 20 reads with two species, A and B, whose abundance ratio is 1:1 and genome length ratio is 1:1. If 6 reads are unique mapped to species A, 4 reads are unique mapped to species B, and the remaining 10 reads are mapped to both species A and species B, then according to the statistical model of Centrifuge, in which the value of *C*_*ij*_ is only 1 or 0 according to whether read *i* is mapped to species *j*, the estimated abundances of species A and species B are 0.6 and 0.4 respectively. When the influence of reference genome similarity is considered in the statistical model of Centrifuge+ by introducing the unique mapping rate, the estimated abundances of species A and species B are 0.57 and 0.43 respectively and more closer to true abundances. Therefore, Centrifuge+ can improve the accuracy of abundance estimates than Centrifuge according to the above discussion.

## 4 Conclusion

Because Centrifuge can analyze not only short reads, but also long reads, Centrifuge has a wide range of application scenarios, such as Pavian (Breitwieser and Salzberg, 2020) and minoTour (Munro *et al*., 2022). Centrifuge is especially applied for ONT shotgun sequencing analysis and is now included as a step in WIMP, which is a quantitative analysis tool for real-time species identification based on the MinIon released by Oxford Nanopore Technologies. In contrast to Centrifuge, Centrifuge+ improved the accuracy of abundance estimates by modifying the statistical model in Centrifuge. The more accurate abundance estimates will be benefit to improve the precision-recall analysis for species identification. Hence, Centrifuge+ will be more widely applied for metagenomic analysis, particularly for real-time species identification.

