## Supplemental Figure 1 and Table 1 for "Centrifuge+: improving metagenomic analysis upon Centrifuge"

### 1. The accuracy tests of Centrifuge+ and Centrifuge

We classified the simulated reads from the file bacteria_sim10K.fa (Kim *et al.,* 2016) with Centrifuge+ and Centrifuge (Kim *et al.,* 2016) and estimated the sequence abundance at species level for each program. The used database consists of 4278 prokaryotic genomes (Kim *et al.,* 2016). In order to evaluate the performance of each program on abundance estimation, we compared the true abundance with the estimated abundance of each program on log-scaled by using Pearson correlation analysis. Here, we define the worst abundance estimate as the lowest log-scaled abundance estiamte. Furtherly, we computed recall, precision and F1 score to evaluate the performance of each program on metagenomic classification at species level. Precision is the proportion of true positive species identified in the sample divided by the number of total species identified by the program and recall is defined as the proportion of true positive species divided by the number of distinct species actually in the sample. The F1 score is the harmonic mean of recall and precision, weighting them equally in a single metric. The species that the number of reads is at least forty is considered as actual species in the sample when computing recall.

### 2. Running Centrifuge and Centrifuge+

The difference between Centrifuge+ and Centrifuge lies in the statistical model for abundance estimation. Therefore, the running mode of Centrifuge+ is the same with Centrifuge. The following is how to run Centrifuge.

perl /path/to/centrifuge-compress.pl B_only taxonomy/ -map seqid_to_taxid.map –o centrifuge_db

/path/to/centrifuge-build –p 4 –conversion-table centrifuge_db.map –taxonomy-tree taxonomy/nodes.dmp –name-table taxonomy/names.dmp centrifuge_db.fa centrifuge_db

/path/to/centrifuge --no-abundance -f -x centrifuge_db –U bacteria_sim10K.fa

### 3. Simulation Results

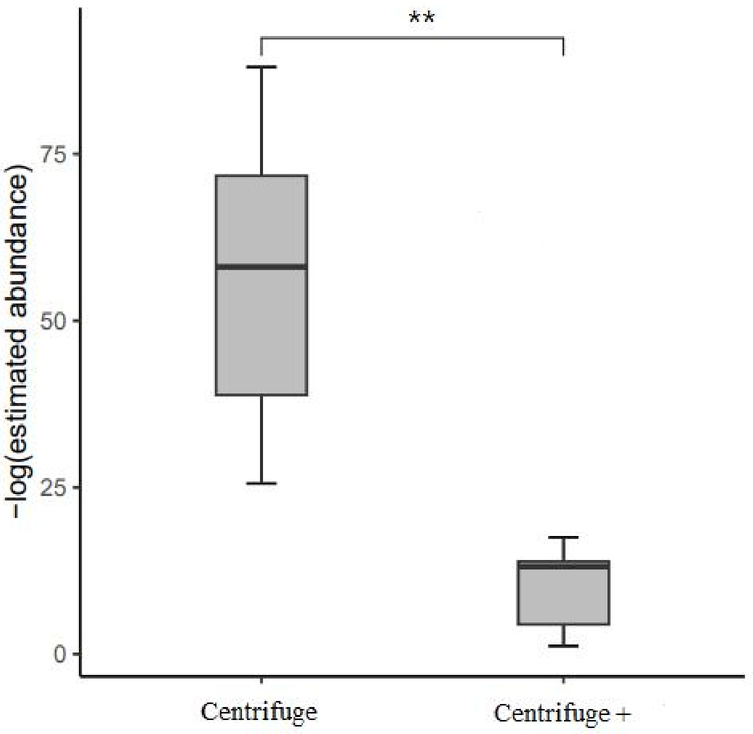


**Fig. S1.** The distribution of the five percent worst abundance estimates on negative log-scaled. The abundance estimation of Centrifuge+ is significantly better than the of Centrifuge. The '**' indicates significant difference of $p$-value < 0.01.

**Table S1.** Classification recall, precision and F1 score for Centrifuge+ and Centrifuge using simulated reads.

| | recall | precision | F1 score |
|---|---|---|---|
| Centrifuge+ | 75.86% | 98.33% | 0.86 |
| Centrifuge | 65.51% | 100% | 0.79 |

Note. The classification recall, precision and F1 score are at species level.
